# Miltefosine increases macrophage cholesterol release and inhibits NLRP3-inflammasome assembly and IL-1β release

**DOI:** 10.1101/430769

**Authors:** Amanda J Iacano, Harvey Lewis, Jennie E Hazen, Heather Andro, Jonathan D Smith, Kailash Gulshan

## Abstract

Miltefosine is an FDA approved oral drug for treating cutaneous and visceral leishmaniasis. Leishmania is a flagellated protozoa, which infects and differentiates in macrophages. Here, we studied the effects of Miltefosine on macrophage’s lipid homeostasis, autophagy, and NLRP3 inflammasome assembly/activity. Miltefosine treatment conferred multiple effects on macrophage lipid homeostasis leading to increased cholesterol release from cells, increased lipid-raft disruption, decreased phosphatidylserine (PS) flip from the cell-surface, and redistribution of phosphatidylinositol 4,5-bisphosphate (PIP2) from the plasma membrane to actin rich regions in the cells. Enhanced basal autophagy, lipophagy and mitophagy was observed in cells treated with Miltefosine vs. control. Miltefosine treated cells showed marked increased in phosphorylation of kinases involved in autophagy induction such as; Adenosine monophosphate-activated protein kinase (AMPK) and Unc-51 like autophagy activating kinase (ULK1). The Toll like receptor (TLR) signaling pathway was blunted by Miltefosine treatment, resulting in decreased TLR4 recruitment to cell-surface and ~75% reduction in LPS induced pro-IL-1β mRNA levels. Miltefosine reduced endotoxin-mediated mitochondrial reactive oxygen species and protected the mitochondrial membrane potential. Miltefosine treatment induced mitophagy and dampened NLRP3 inflammasome assembly. Collectively, our data shows that Miltefosine induced ABCA1 mediated cholesterol release, induced AMPK phosphorylation and mitophagy, while dampening NLRP3 inflammasome assembly and IL-1β release.

**Significance Statement:** Atherosclerosis is driven by cholesterol accumulation and inflammation, and the arterial macrophage is a key cell type in both of these processes. The macrophage characteristics that protect against atherosclerosis include increased cholesterol efflux/reverse cholesterol transport, increased autophagy, and deceased inflammatory cytokine production and signaling. Here, we show that one single orally available compound, Miltefosine, can target multiple macrophage pathways involved in lipid homeostasis and inflammation. Miltefosine activated cholesterol release and autophagy while inhibiting pro IL-1β gene expression and NLRP3 inflammasome assembly. Miltefosine activated AMPK signaling pathway and mitophagy, leading to reduced NLRP3 inflammasome assembly and IL-1β release.

## Introduction

Atherosclerosis, a sterile inflammatory disease, is the major cause of coronary artery disease (CAD). LDL cholesterol drives atherosclerosis by depositing LDL into the arterial intima, where it can be modified to induce endothelial cell activation and the recruitment of leukocytes; and, the uptake of modified LDL into macrophages leads to foam cell formation and a further amplification of inflammation^1–4^. Accumulation of oxidized lipids and cholesterol crystals in plaques activate toll-like receptor (TLR) pathways and the assembly of the NLRP3 inflammasome^2,5^. The NLRP3 inflammasome plays a key role in processing procaspase 1 resulting in subsequent caspase 1 mediated processing of pro IL-1β to generate active interleukin-1β (IL-1β). The role of IL-1β in promoting human CAD was highlighted by the recently concluded CANTOS trial, showing that anti-IL-1β therapy met the primary endpoint, a reduction in a composite of heart attack, stroke and cardiovascular death^6^.

In contrast to atherogenic pathways, the atheroprotective pathways such as autophagy and cholesterol efflux become increasingly dysfunctional in aging, advanced atherosclerotic plaques, and in animal models of atherosclerosis and diabetes^5,7-9^. Thus, the simultaneous induction of atheroprotective pathways, along with dampening of atherogenic pathways may serve as a potent therapeutic treatment for CAD patients. Here, we report that Miltefosine, an FDA approved drug for treating visceral and cutaneous leishmaniasis, promoted cholesterol release, disrupted lipid-rafts and TLR4 signaling, increased cell-surface phosphatidylserine (PS) exposure, induced phosphatidylinositol 4,5-bisphosphate (PIP2) trafficking from plasma membrane (PM) to the cell interior. Miltefosine treated cells exhibited increased phosphorylation of autophagy inducing kinases AMPK1 and ULK1, leading to increased basal autophagy, lipophagy and mitophagy. The lipopolysaccharide (LPS) induced mitochondrial reactive oxygen species (ROS) generation and NLRP3 inflammasome assembly was blunted in cells pretreated with Miltefosine, leading to decreased caspase 1 cleavage and mature IL-1β release. The structure of Miltefosine is similar to lyso-PC and it is known to integrate in the cell membrane and redistribute in intracellular membranes of ER, Golgi and mitochondria^10^. It is widely believed that most of the downstream effects of Miltefosine are dependent on cell-type, with a range of activities such as anticancer, antimicrobial, effects on cholesterol homeostasis, and inhibition of Mast cell activation^10–13^. The detailed investigation of mechanisms involved in Miltefosine’ s action may lead to novel therapeutic targets for preventing and treating cardiovascular and inflammatory metabolic diseases.

## Results

### Miltefosine induced ABCA1 mediated cholesterol release from cells

Previous study have shown that Miltefosine and other alkylphospholipids induced cholesterol efflux from the HepG2 cells^14^. We tested effect of Miltefosine treatment on ABCA1 mediated cholesterol release in RAW264.7 macrophages. Control and ABCA1 expressing cells were incubated with different doses of Miltefosine for 4h and cholesterol release to serum-free media in the absence of apoA1, the usual acceptor, was determined. As shown in ***Fig. 1A***, Miltefosine treatment of control cells without ABCA1 expression led to a moderate, but significant, increase in cholesterol release to media with 7.5 μM showing ~1 % increase (n=4, p<0.001) vs. control cells. The ABCA1-expressing cells showed more robust cholesterol release to acceptor-free media in presence of Miltefosine with 7.5 μM showing ~5.28 % increase (n=4, p<0.01) vs. control cells. Miltefosine can increase ABCA1 mediated cholesterol release by increasing either total or cell-surface levels of ABCA1. We found no significant increase in ABCA1 levels (***Fig. 1B***), instead a trend toward decreased ABCA1 levels was observed in Miltefosine treated cells. This data indicated that the Miltefosine mediated increase in cholesterol release was not due to increased ABCA1 levels. Further studies used 7.5 μM or less of Miltefosine, as these doses did not lead to cell death in RAW264.7 macrophages or in mouse bone-marrow derived macrophages (BMDMs) (***Fig. S1A, S1B***). We observed the Miltefosine induction of cholesterol release in three additional cell lines, HEK293, BHK cells and THP-1 macrophages. In all cases, addition of Miltefosine induced basal cholesterol release, and markedly induced ABCA1-mediated cholesterol release to the acceptor-free media (***Fig. 1C, S1C, S1D***). Addition of cholesterol acceptor apoA1 to Miltefosine treated cells did not lead to further significant increase in cholesterol release (***Fig. S1D, S1E, S1F***). To test if the lipid floppase activity (outward translocation across the plasma membrane) of ABCA1 is required for increased cholesterol release in Miltefosine treated cells, we used HEK293 cells stably expressing WT-ABCA1 isoform or a Tangier disease double mutant W590S-C1477R-ABCA1 isoform. The WT ABCA1 can flop phosphatidylserine (PS) and phosphatidylinositol 4,5-bisphopshate (PIP2) across plasma membrane^15^. The W590S mutation in ABCA1 decreases PS flop^16^ while C1477R decreases PIP2 flop^15^. The W590S-C1477R double mutant (DM) isoform of ABCA1, with protein expression levels slightly higher than WT ABCA1 (***Fig. S1G***), showed significantly lower cholesterol release to acceptor-free media compared to WT-ABCA1 (***Fig. 1D***), indicating that lipid floppase activity of ABCA1 promotes the Miltefosine induced cholesterol release to media. It has previously been shown that ABCA1 expression promotes membrane blebbing and the release of microparticles^17,18 19^. Microparticles are generated by ABCA1 activity, but unlike nascent HDL particles, the microparticles are formed even in the absence of apoA1^17^. We determined the effect of Miltefosine on generation of microparticles by cells. The media from RAW264.7 macrophages incubated ± Miltefosine for 4h was collected and analyzed by a small particle tracking analyzer. The media from the cells treated with Miltefosine had significant 2-fold more microparticles compared to control cells (P<0.001, ***Fig. S2A***). These data indicated that Miltefosine mediated remodeling of the plasma membrane promotes generation of microparticles leading to cellular cholesterol release to the media, which was even greater in the presence of ABCA1 expression.

**Figure 1:**
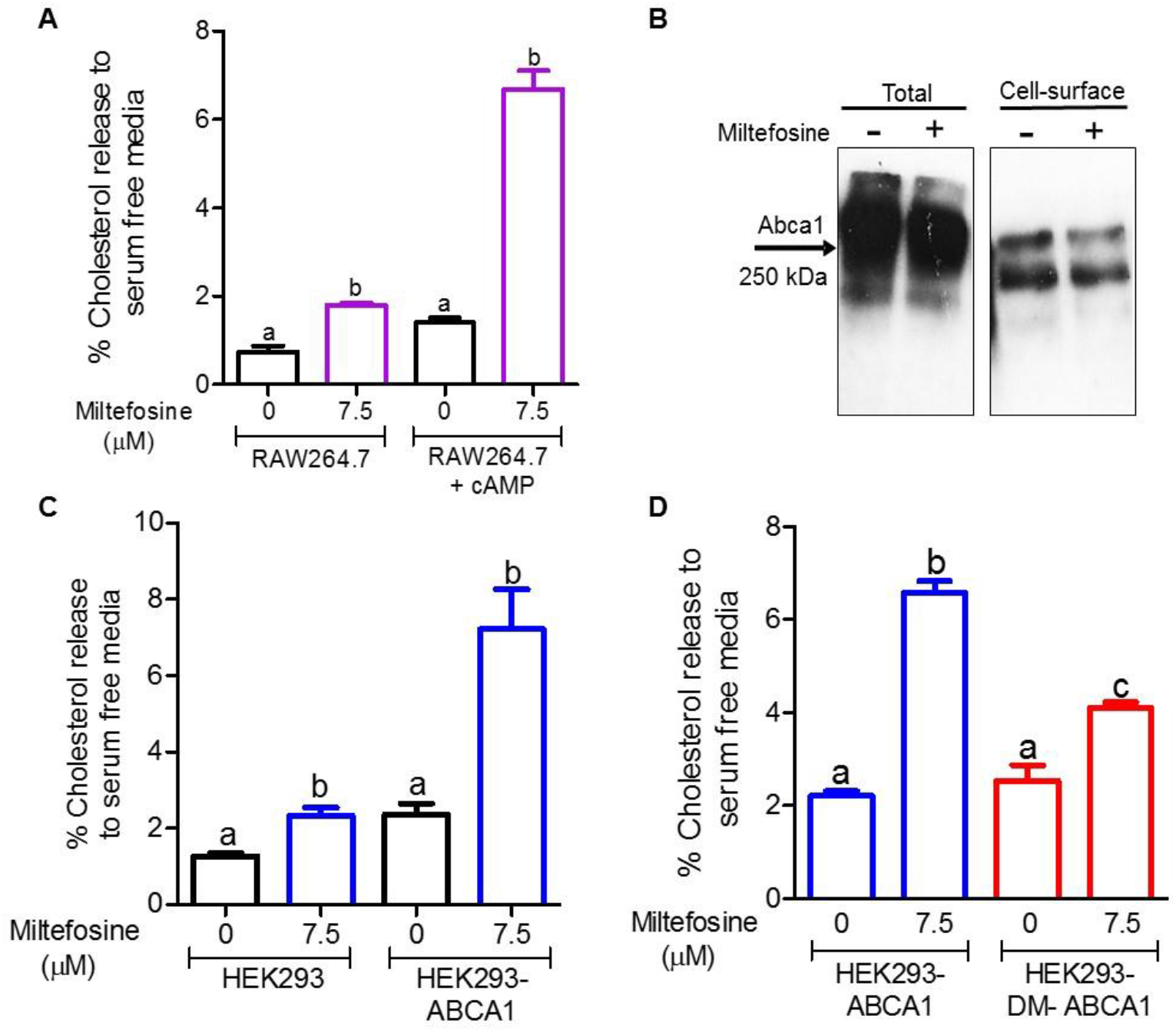
Miltefosine increases ABCA1 mediated cholesterol release. ***A)*** RAW264.7 macrophages were labeled with ^3^H-cholesterol and pretreated with or without 300 μM 8Br-cAMP to induce ABCA1 expression. Cholesterol release to chase media was performed for 4h at 37°C in serum-free DMEM without addition of acceptor containing either vehicle or 7.5 μM Miltefosine. Values are % cholesterol release mean ± SD, N=5, different letters above the bars show p<0.01 by ANOVA Bonferroni posttest, with separate analyses for ± ABCA1 induction. ***B)*** Western blot analysis of 8Br-cAMP induced RAW264.7 cell total and cell-surface ABCA1 ± 7.5 μM Miltefosine treatment for 4h. ***C)*** Cholesterol release (4h at 37°C) from ABCA1 stably transfected and control HEK293 to serum-free DMEM without addition of acceptor containing either vehicle or 7.5 μM Miltefosine. Values are % cholesterol release mean ± SD, N=3-5, different letters above the bars show p<0.01 by ANOVA Bonferroni posttest, with separate analyses for ± ABCA1 induction. ***D)*** Cholesterol release (4h at 37°C) from ABCA1 and ABCA1 (W590S, C1477R) double mutant (DM) stably transfected HEK293 to serum-free DMEM ± 7.5 μM Miltefosine. Values are % cholesterol release mean ± SD, N=3, different letters above the bars show p<0.01 by ANOVA Bonferroni posttest.

### Miltefosine disrupts lipid rafts and increased PS exposure by inhibiting PS flip

Lipid rafts are comprised mainly of PC, cholesterol and sphingomyelin^20^, and play a role in AKT and TLR signaling pathways. Disruption of lipid-rafts by ABCA1 is proposed to provide a free cholesterol pool for efflux and hamper TLR signaling via reduced recruitment of TLR2/4 to lipid-rafts^21–23^. Previous studies have also shown that Miltefosine uses lipid-rafts as entry portals to cells^24^ and can act as a lipid-raft disrupting agent to inhibit human mast cell activation^13^. We determined the status of AKT phosphorylation in RAW264.7 macrophages and found that Miltefosine potently inhibited basal AKT phosphorylation (***Fig. S2B, S2C***). The lipid-rafts in Miltefosine treated RAW264.7 macrophages were probed by staining for the ganglioside GM1 using Alexa647-labeled cholera toxin B, followed by either fluorescent microscopy or flow cytometry. As shown in ***Fig. 2A*** cells treated with 7.5 μM Miltefosine for 16h displayed less GM1 vs. control cells. Flow cytometry quantification showed ~ 26% GM1 decrease in Miltefosine treated vs. control cells (n=3, p<0.004 by t-test, ***Fig. 2B***). We have previously shown that membrane remodeling by sphingomyelin depletion resulted in lipid raft disruption, increased cell-surface PS, and promoted cholesterol extraction from cells^25^. A flowcytometry based Annexin Cy5 binding assay was used to investigate if Miltefosine increased PS exposure at the cell-surface. Cells expressing ABCA1 were used as positive control, as PS cell-surface exposure is increased by ABCA1’s PS flop activity^15,16^. The Miltefosine treated cells showed ~24% increase in cell surface PS vs. untreated cells (n=4, p<0.005) (***Fig. 2C)***. As expected, the cells expressing ABCA1 also showed increased PS exposure (~23% increase vs control cells, n=4, p<0.05). Cell surface PS was further increased by Miltefosine treatment in ABCA1 expressing cells (~39% increase vs. control cells, n=4, p<0.0005) ***(Fig. 2C, S2D)***. These data indicates that Miltefosine increased cell-surface PS exposure independent of ABCA1 PS floppase activity. The increased levels of cell surface PS may be due to the decreased rate of PS flipping (inward translocation). To qualitatively observe the PS flipping across plasma membrane (PM), RAW264.7 cells ± 7.5uM Miltefosine treatment for 16h, were incubated with exogenously added NBD-PS at final concentration of 25 μM. In control cells, the majority of exogenous NBD-PS moved to intracellular compartments over a 15 min incubation period at 37°C, while Miltefosine treated cells displayed more NBD-PS at the PM and less inside the cells (***Fig. 2D***). To quantitatively measure flipping of NBD-PS, we used a previously described flow-cytometry based method using a membrane-impermeable NBD quenching reagent sodium dithionite^25^. As shown in ***Fig. 2E***, Miltefosine treated cells flipped significantly lower amounts of NBD-PS vs. control cells, ~44% vs. ~70% respectively (n=4, p<0.0002). These data indicate that Miltefosine inhibited PS flip from the cell surface, resulting in increased PS exposure at the cell surface.

**Figure 2.**
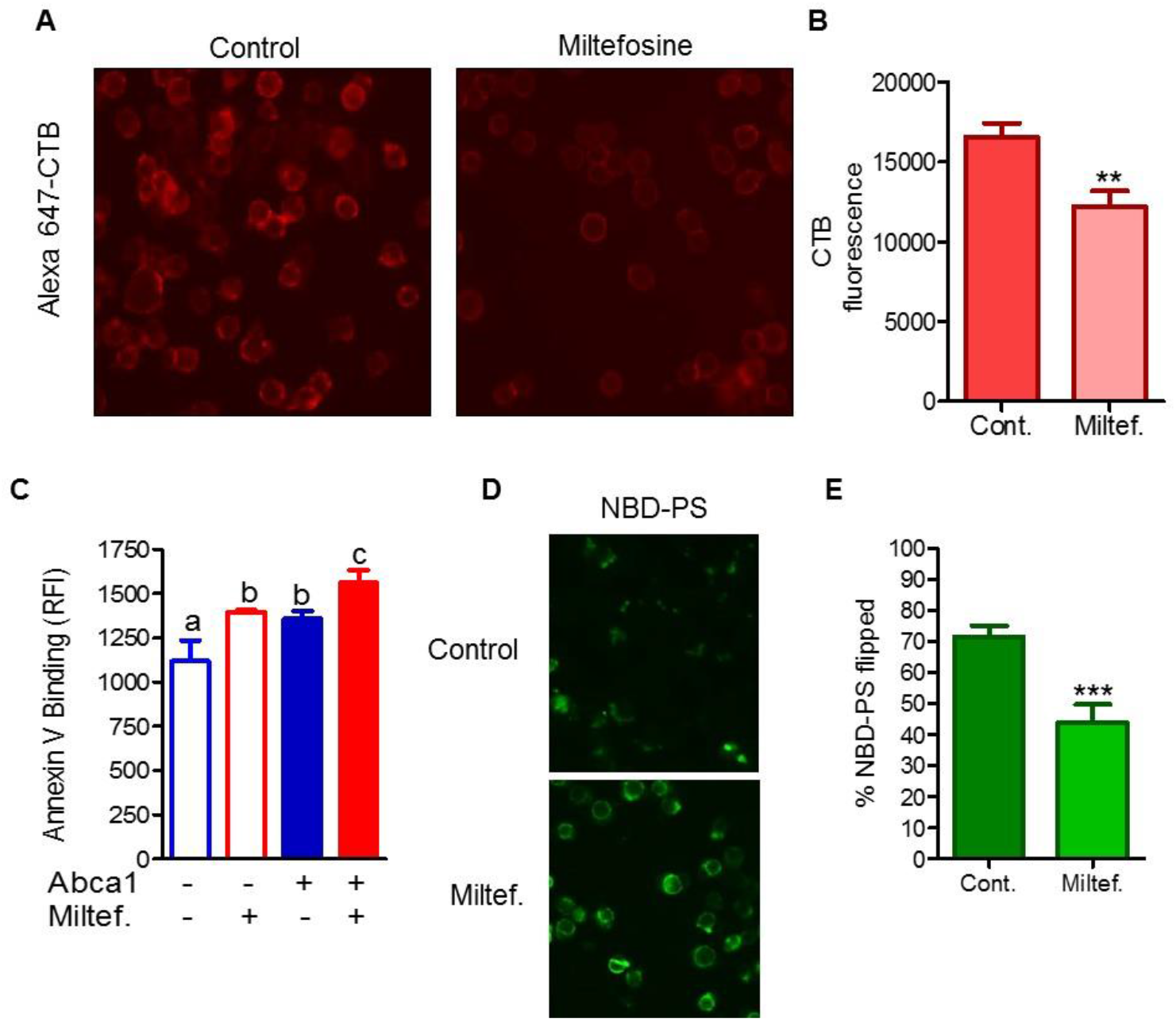
Miltefosine disrupts lipid-rafts and inhibits PS flip across plasma membrane. **A)** GM1 levels assessed by binding of cholera toxin B (CTB) in live RAW macrophages ± 7.5 μM Miltefosine for 16h at 37°C. ***B)*** Flow c_o_ytometry quantification of CTB binding of RAW cells treated ± 7.5 μM Miltefosine for 16h at 37°C. Values are the mean ± SD of the median fluorescence from 3 independent wells (**, p<0.01 by two-tailed t-test). ***C)*** RAW macrophages were incubated with or without 8Br-cAMP to induce ABCA1 and ± 7.5 μM Miltefosine for 16 hrs. PS exposure was determined Annexin V binding via flow cytometry (different letters above the bars show p<0.01 by ANOVA Bonferroni posttest). ***D)*** Cells were pretreated ± 7.5 μM Miltefosine and incubated with 25 μM NBD-PS at RT for 15 min to assess cellular association of PS. ***E)*** Quantification of NBD-PS translocated inside the cells. RAW macrophages were pretreated ± 7.5 μM Miltefosine and incubated with 25 μM NBD-PS at 37°C for 15 min in phenol red free DMEM. The cells were subjected to flow cytometry analysis ± sodium dithionite to quench extracellular NBD fluorescence, yielding only intracellular NBD fluorescence (mean ± SD % NBD-PS translocated into the cells; N = 3; ***, p<0.001 by two-tailed t-test).

### PIP2 is redistributed from plasma membrane upon Miltefosine treatment

Similar to PS, phosphatidylinositol 4, 5-bisphosphate (PIP2) is also sequestered in the inner leaflet of the plasma membrane. To determine the effect of Miltefosine on PIP2 trafficking and localization, RAW264.7 cells stably transfected with a PIP2 reporter plasmid containing 2 copies of the pleckstrin homology domain from phospholipase Cδ fused to GFP (2X-PH-PLC-eGFP) were treated with ± 7.5 μM Miltefosine for 16h and PIP2 reporter localization was determined by fluorescent microscopy. As shown in ***Fig. 3A***, control cells showed uniform localization of the PIP2 reporter at the plasma membrane while the cells treated with Miltefosine showed an additional localization of the reporter in a specific actin rich region of the cytoplasm in both fixed and live cells (***Fig. 3B, C, S3***).

**Figure. 3.**
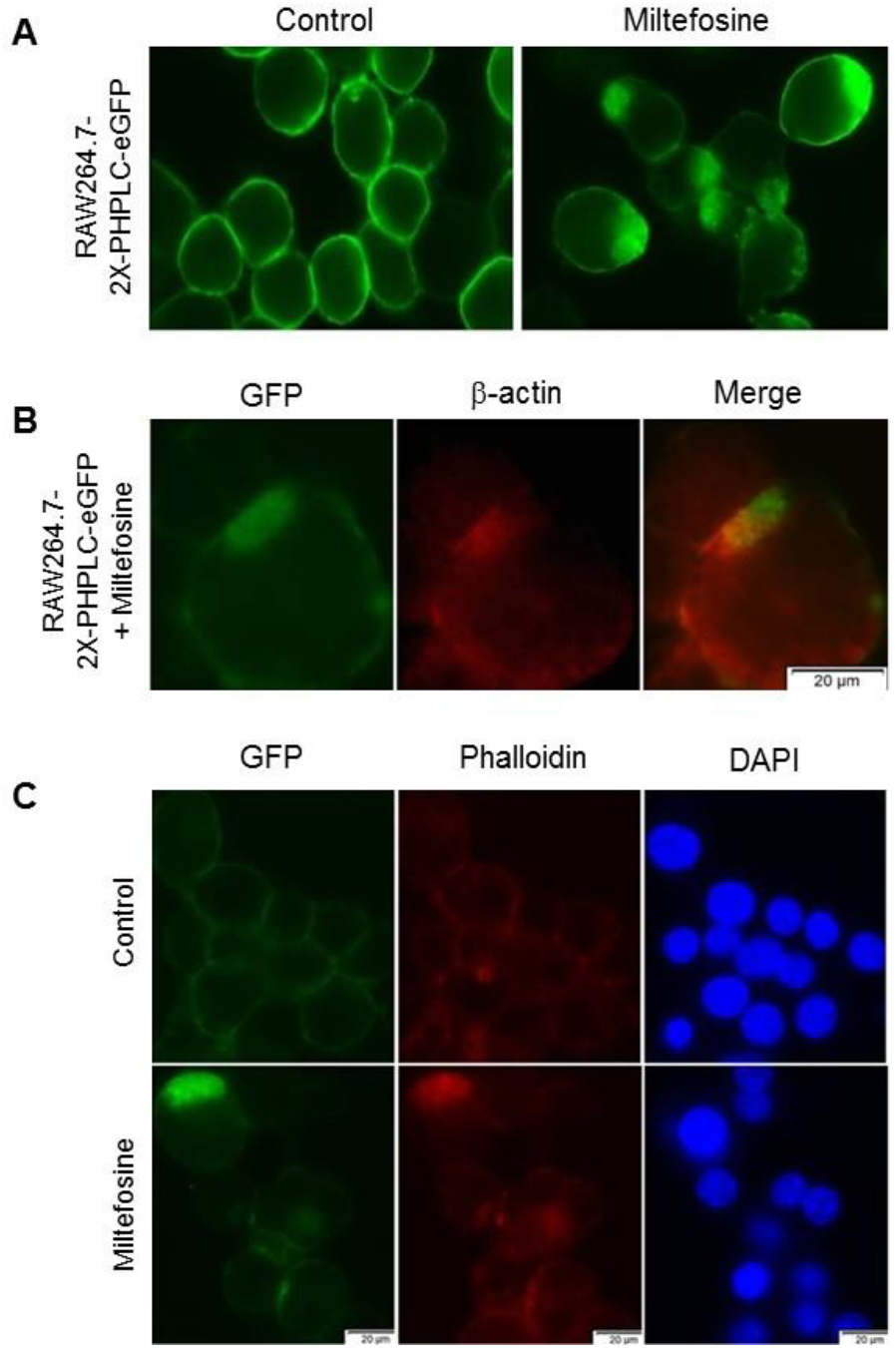
Miltefosine alters plasma membrane PIP2 localization. ***A***) RAW264.7 cells were stably transfected with 2X-PH-PLCeGFP reporter plasmid and treated ± 7.5 μM Miltefosine for 12h at 37°C and the localization of the GFP-tagged PIP2 reporter was observed. ***B***) RAW264.7 cells treated as in (A) were fixed, permeabilized, and stained with mouse anti β-actin antibody to observe colocalization of actin with the PIP2 reporter. ***C)*** RAW264.7 cells stably transfected with 2X-PH-PLCeGFP reporter plasmid ± 7.5 μM Miltefosine treatment, stained for β-actin with Phalloidin and DAPI counterstained.

### Miltefosine induced basal and lipid-droplet autophagy

Autophagy plays a protective role against atherosclerosis^8^ and becomes increasingly impaired in advanced human plaques^7^. Autophagy is required for hydrolyzing macrophage cholesterol ester stored in lipid droplets to generate free cholesterol that can be effluxed from cells^8^. As Miltefosine increases cholesterol release, we tested if Miltefosine induced autophagy. We observed that the RAW264.7 macrophages treated with 7.5 μM Miltefosine for 16h showed multiple p62 puncta in cytoplasm, while control cells had evenly distributed weak cytoplasmic signal with almost no p62 puncta signal (***Fig. 4A***, upper panel). Miltefosine treatment also increased LC3 puncta formation in macrophages transfected with LC3-GFP plasmid (***Fig. 4A*** lower panel). Next, we evaluated whether the LC3 puncta formation was due to increased autophagic initiation or due to decreased autophagic flux. Decreased or blocked autophagy flux can also lead to accumulation of autophagic markers. If western blot analysis shows increased LC3-II after chloroquine (an autophagy flux inhibitor) treatment, then autophagic flux is occurring. We found that the LC3-II protein levels were increased by 3.5-fold in Miltefosine treatment alone vs. control cells (***Fig. 4B, 4C, S4***), while there was an additional 1.8 fold increase in LC3-II protein levels with Miltefosine + chloroquine treatment compared to cells treated with Miltefosine alone (***Fig. 4C***). These data indicated that Miltefosine increased autophagic flux. Next, we tested the effect of Miltefosine on generation of free cholesterol from cholesterol ester rich lipid droplets. RAW264.7 macrophages were loaded with 100 μg/ml acetylated LDL (AcLDL) for 24h followed by a 4h chase with an acyl-co A cholesterol O-acyltransferase (ACAT) inhibitor (ACATi) to prevent cholesterol re-esterification ± 7.5 μM Miltefosine. The total and free cholesterol levels were determined by enzymatic assay described previously^26^. As expected, the AcLDL loaded cells had a higher ratio of cholesterol ester to free cholesterol. Miltefosine treatment led to decreased CE: FC ratio, indicating that Miltefosine treated cell had higher free cholesterol and less cholesterol esters as compared to control cell (P<0.05 vs. control) (***Fig. 4D***). The cells chased with ACATi had reduced CE:FC ratio (P<0.001 vs. control), while the Miltefosine + ACATi treated cells showed further lowering of CE:FC ratio (p<0.001 vs. control) (***Fig. 4D***). We also tested the effect of Miltefosine on lipid-droplets by loading RAW264.7 cells with 100μg/ml AcLDL for 24h, followed by a 4 h chase with ± Miltefosine and staining with Nile-red dye. Cells treated with Miltefosine showed reduced Nile-red staining as compared to control cells (***Fig. S4D***). To quantify the lipid-droplets, AcLDL loaded cells were chased for 4h with ACATi ± Miltefosine and subjected to flow-cytometry analysis. A shown in ***Fig. 4E***, the cells chased with ACATi showed decreased lipid droplets that were further decreased by Miltefosine (n=4, p<0.0001). These data indicate that Miltefosine induced lipid-droplet autophagy as ACATi prevents new CE formation.

**Figure. 4.**
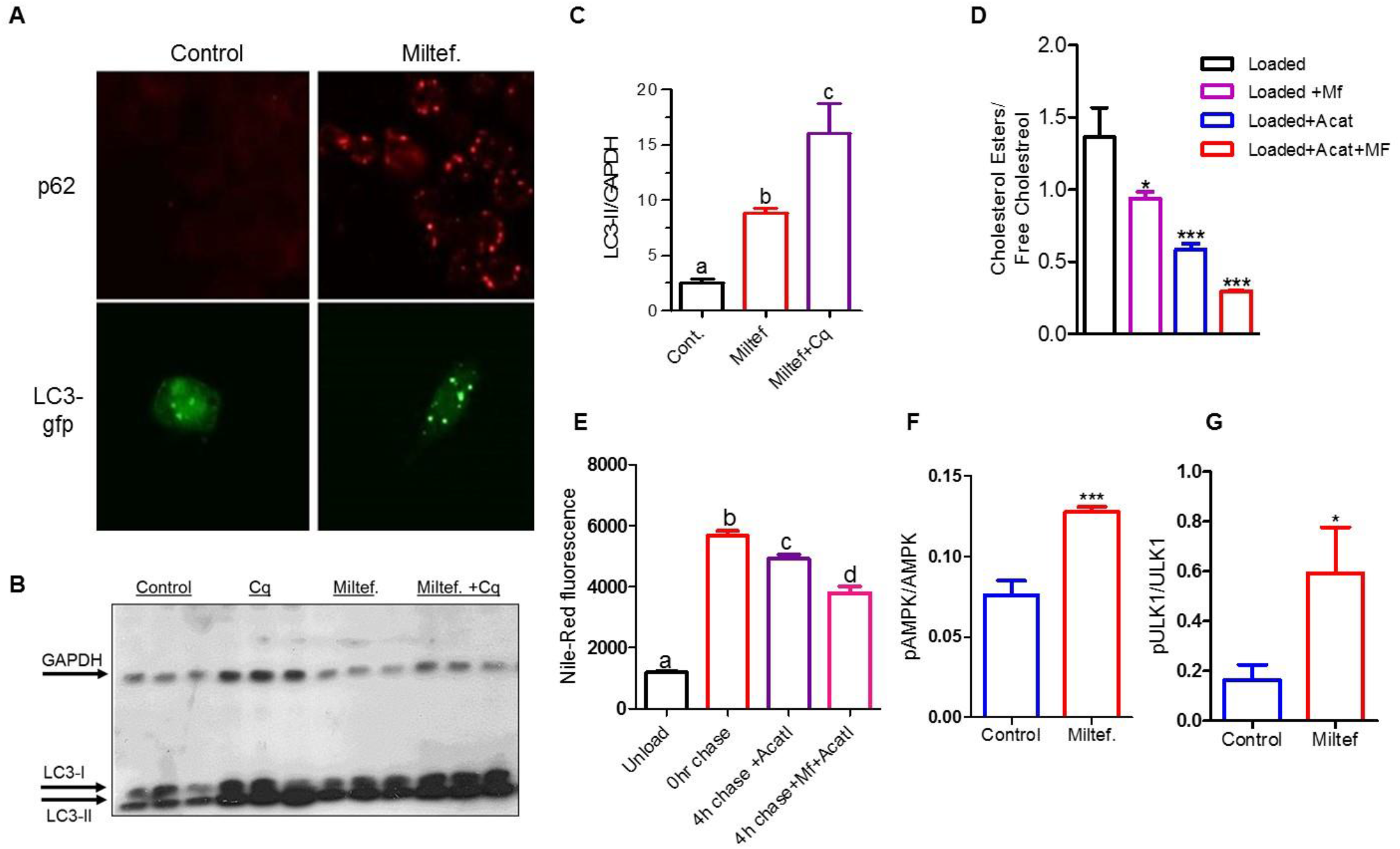
Miltefosine induces autophagy. ***A)*** RAW264.7 cells or LC3-GFP transfected RAW264.7 cells (lower panel) were treated with ± 7.5 μM Miltefosine, followed by staining with p62 antibody (red, upper panel). LC3-GFP protein (green) localization is shown in the lower panel. ***B, C)*** RAW264.7 cells were treated ± 7.5 μM Miltefosine and autophagic flux was assessed in presence of 30 μM chloroquine for the last 2 h of incubation. The amount of LC3-II was assessed by western blot and densitometry analysis relative to GAPDH (mean ± SD, N = 3, different letters above the bars represent p<0.001 by ANOVA with Bonferroni posttest). ***D)*** Cellular CE/FC ratio in RAW264.7 cells loaded with 100 μg/ml acetylated low-density lipoprotein (AcLDL) and addition of 2 μg/ml ACAT inhibitor (ACATi) and/or 7.5 μM Miltefosine as indicated (N=3, mean ± SD; *, p<0.05 vs. control; ***, p<0.001 vs. control by ANOVA Bonferroni posttest). ***E)*** Cellular neutral lipids assayed by Nile red staining and flow cytometry ± AcLDL loading, ± 4 h chase ± 7.5 μM Miltefosine ±2 μg/ml ACATi, as indicated (mean ± SD, N = 4, different letters above the bars represent p<0.0001 by ANOVA with Bonferroni posttest). The densitometry analysis of p-AMPK (***F***) or p-ULK1 (***G***) relative to AMPK or ULK1 (mean ± SD, N = 3; ***, p<0.001, *, p<0.05, by two-tailed t-test).

Autophagy induction is promoted by energy sensing Adenosine monophosphate-activated protein kinase (AMPK), while mammalian target of rapamycin (mTOR) acts to inhibit autophagy. The AMPK substrate ‘Unc-51 like autophagy activating kinase (ULK1)’ is also phosphorylated by AMPK during autophagy and both AMPK1 and ULK1 have been shown to be essential for mitophagy and cell survival during starvation^27 28,29^. In order to determine the mechanism of autophagy induction by Miltefosine, we tested the phosphorylation status of AMPK and ULK1 and found that the phosphorylation levels of both AMPK and ULK1 were significantly higher in Miltefosine treated cells vs. control cells (***Fig. 4F, 4G, S4E, S4F***). These data indicated that Miltefosine induced autophagy and did not block cellular autophagy flux.

### Miltefosine decreased LPS mediated induction in IL-1β levels

We demonstrated in ***Fig 2A*** that Miltefosine reduces lipid-rafts. The lipid rafts have been shown to play an essential role in TLR2/4 signaling pathway and atherosclerosis progression^21,22,30–32^. We tested if LPS mediated recruitment of TLR4 to the cell surface is impaired in Miltefosine treated cells. The BMDMs ± pretreatment with Miltefosine for 16h were analyzed for TLR4 cell-surface recruitment by flow cytometry. As expected, the LPS treated control cells showed increased binding of TLR4 antibody, indicating robust TLR4 recruitment to the cell-surface. In contrast, the cells pretreated with Miltefosine showed significantly reduced TLR4 recruitment to cell-surface (***Fig. 5A, S5A, S5B***). Next, we tested if LPS mediated induction of pro IL-1β mRNA was affected by Miltefosine. Bone-marrow derived macrophages (BMDMs) ± Miltefosine pretreatment were incubated with 1 μg/ml LPS for 4h, followed by qRT-PCR using IL-1β specific primers and control Actb (β-actin) primers. The LPS induction of IL-1β mRNA was markedly reduced in Miltefosine treated cells (~4.5 fold decrease, n=3, p<0.034) (***Fig. 5B***). Western blot analysis of pro IL-1β protein also revealed a similar effect of Miltefosine with a ~55% decrease in protein levels (***Fig. 5C***). Taken together, these data indicated that Miltefosine disrupted lipid-rafts, decreased LPS mediated TLR4 cell-surface recruitment and induction of IL-1 β mRNA and protein levels in macrophages.

**Figure. 5.**
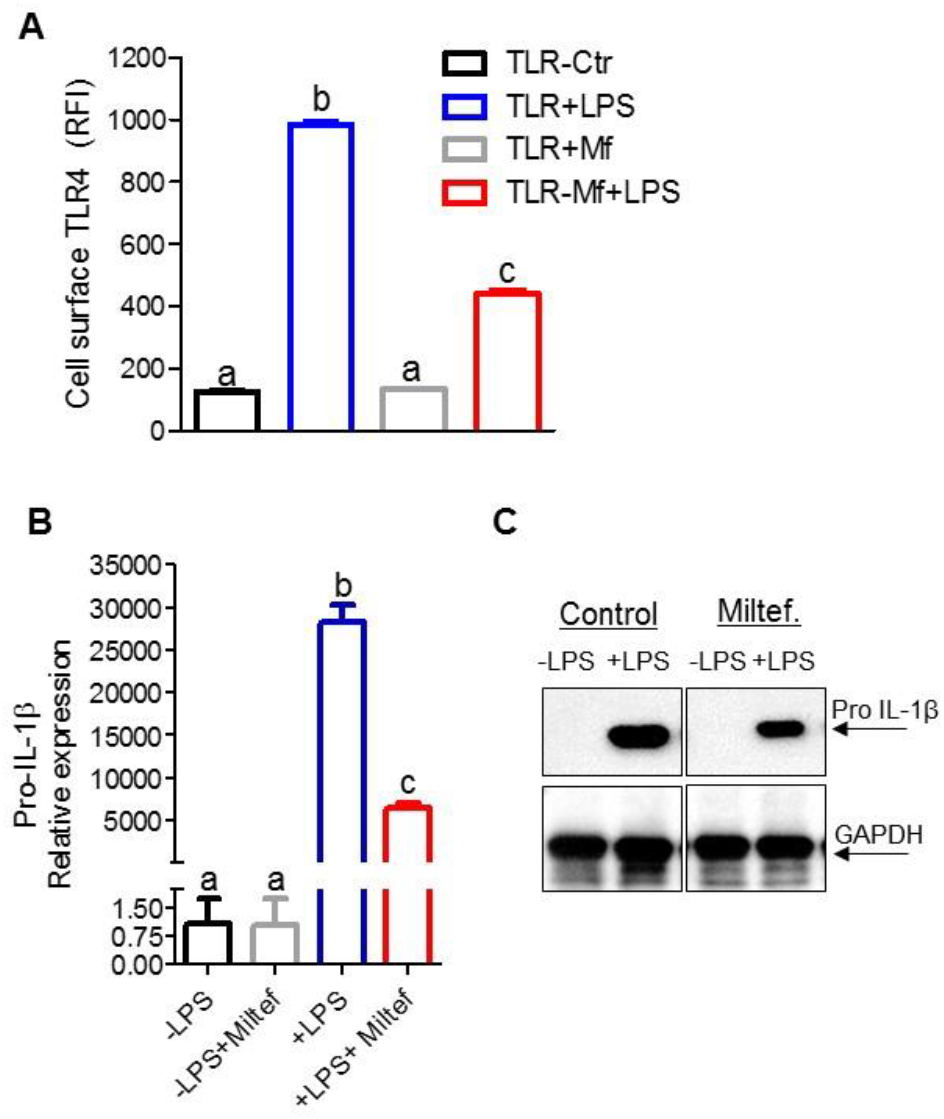
Miltefosine dampens TLR4 signaling in bone marrow derived macrophages. ***A)*** Mouse BMDMs were treated with ± 5 μM Miltefosine for 16h at 37°C and primed by incubation with ± 1μg/ml LPS at 37°C for 4hrs, followed by analysis of TLR4 antibody binding by flow cytometry (N=3, different letters above the bars show p<0.01 by ANOVA Bonferroni posttest. ***B)*** Mouse BMDMs were treated as in (A), followed by analysis of pro-IL-1β mRNA with β-actin mRNA used as a control (N=6, mean ± SD; different letters above the bars represent p<0.01 by ANOVA with Bonferroni posttest). ***B)*** Mouse BMDMs were treated with ± 5 μM Miltefosine and primed by incubation with ± 1mg/ml LPS followed by western blot analysis for IL-1 β with GAPDH used as loading control.

### Miltefosine inhibited NLRP3 inflammasome assembly and IL-1β release

Cholesterol crystals can activate NLRP3 inflammasome and promote atherosclerosis^2^. Cholesterol crystals and other treatments including extracellular ATP can promote NLRP3 inflammasome assembly, which processes pro IL-1 β to its mature form for secretion from cells^2,33^. We tested if Miltefosine effected NLRP3 inflammasome assembly. The BMDMs pretreated with ± 5 μM Miltefosine for 16h, were primed with LPS for 4h and incubated with ATP for 20 min, followed by probing with anti-ASC antibody for visualization of NLRP3 inflammasome assembly. BMDMs treated with LPS and ATP formed ASC specks in 41% cells (n=308 cells analyzed) while in BMDMs pretreated with Miltefosine only 12% of cells showed ASC specks (523 cells analyzed, p<0.0001 by two-tailed Fisher’s exact test) (***Figs.6A, S6, S7***). Miltefosine treatment led to markedly reduced release of cleaved caspase-1 in media and ~75% decrease in release of mature IL-1β to media as compared to control macrophages (n=3, p<0.0039) (***S8A, Fig. 6B***,). To decipher the mechanism by which Miltefosine inhibited NLRP3 inflammasome assembly, we determined the mRNA and protein levels of the constituents of the NLRP3 inflammasome and one substrate Gasdermin D (Gsdmd) in BMDMs with or without LPS priming. Miltefosine with LPS actually slightly increased the mRNA levels, beyond the effects of LPS alone, for NLRP3, procaspase 1 and Gasdermin D (***Fig. 6C***). In agreement with earlier published studies^34^, western blot analysis showed robust induction in protein levels of NLRP3 upon LPS treatment, which was not reduced by Miltefosine treatment (***Fig. S8B)***. Miltefosine did not appreciably alter the protein levels of ASC or procaspase 1 ***(Fig. S8C, S8D)***. To test if Miltefosine specifically inhibited NLRP3 inflammasome assembly and IL-1β release, we determined the effects of Miltefosine on the AIM2 inflammasome. The control and Miltefosine treated macrophages were primed with LPS (1mg/ml) for 4hrs followed by lipofectamine mediated transfection of poly (dA:dT) for 3h. As shown in ***Fig. 6D***, there was no difference in levels of AIM2 induced IL-1 β released from cells + Miltefosine treatment (n=3, n.s.); thus, Miltefosine did not alter AIM2 inflammasome activity. The release of mature IL-1β from cells is regulated by Gasdermin D mediated pore formation on the plasma membrane^35,36^ and previous studies have shown that the deficiency of ABCA1/ABCG1 increases inflammation in macrophages while ABCA1 mediated cholesterol efflux in dendritic cells dampens inflammasome assembly^37,38^. Though, we used unloaded BMDMs for these studies, there is a possibility that Miltefosine inhibits NLRP3 inflammasome assembly via cholesterol depletion. To directly test the role of cholesterol depletion on NLRP3 inflammasome assembly, we treated BMDMs with 5μM Miltefosine for 16h or with 1mM cyclodextrin for 45 minutes at 37°C, followed by LPS/ATP incubation. Although cholesterol levels in BMDMs treated with cyclodextrin were significantly lower compared to Miltefosine treated cells (n=3, p<0.01, ***Fig. 6E***), Miltefosine was much more effective than cyclodextrin in inhibiting inflammasome activity as assayed by IL-1β release (***Fig. 6F)*** and ASC speck formation (***Fig. S6, S9***). Thus, NLRP3 inflammasome inhibition by Miltefosine cannot be attributed solely to cholesterol depletion.

**Figure. 6.**
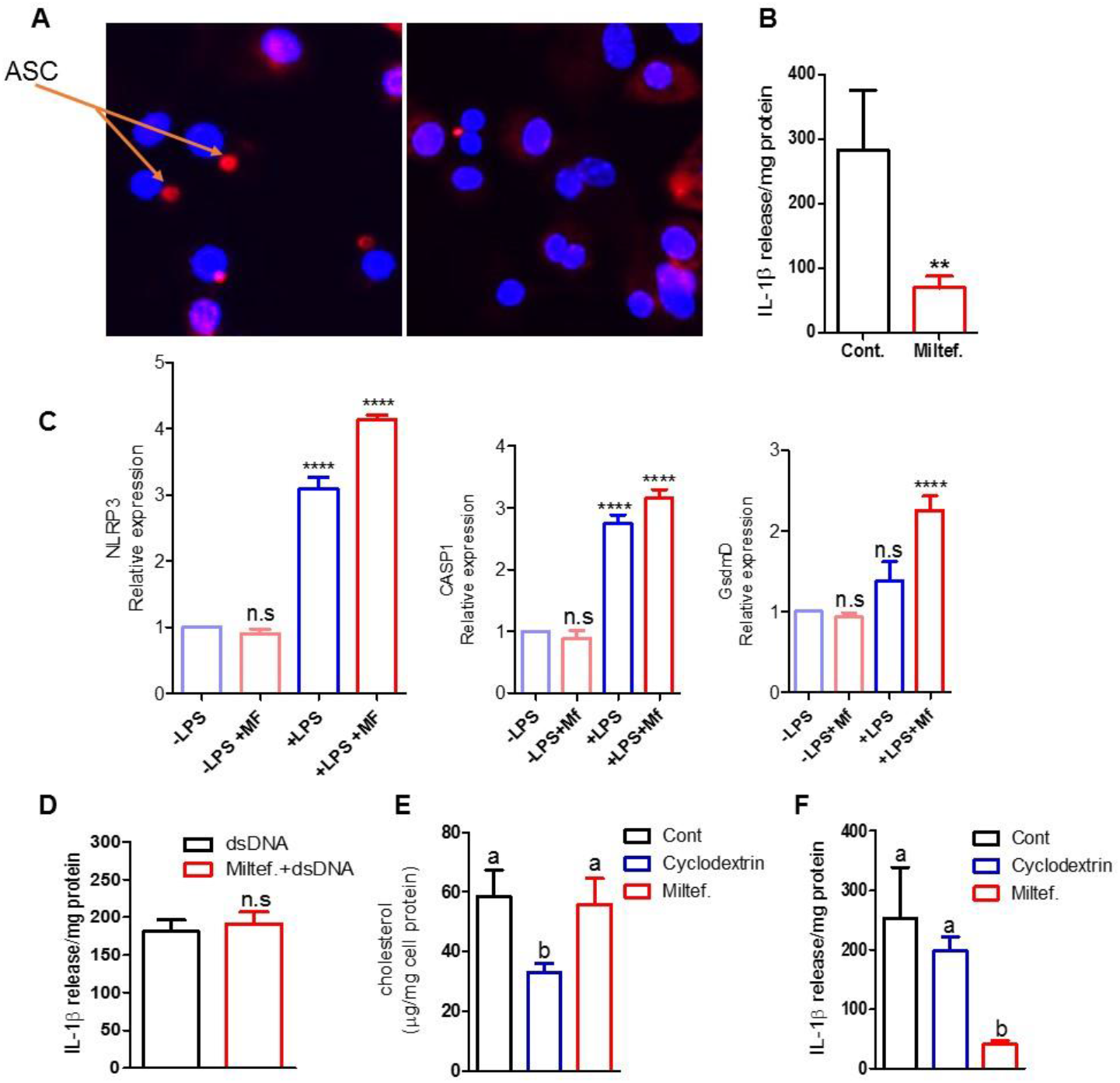
Miltefosine inhibits Nlrp3 inflammasome assembly and IL-1β release. ***A)*** Mouse BMDMs were pretreated with ± 5 μM Miltefosine, primed by incubation with ± 1 μg/ml LPS, and induced for NLRP3 inflammasome assembly by incubation with 1 mM ATP for 20 min. Cells were fixed and stained with anti-ASC with DAPI. ***B)*** IL-1 β levels in media from cells treated with LPS and ATP ± 5 μM Miltefosine pretreatment (N=4, mean ± SD; **p<0.01 by two tailed t-test).***C)*** qRT-PCR analysis of NLRP3, caspase 1, and Gasdermin D (GsdmD) mRNAs, relative to β-actin mRNA, from mouse BMDMs ± 5 μM Miltefosine pretreatment, ± LPS treatment (N=6, mean ± SD; ****p<0.0001 by ANOVA with Bonferroni posttest). ***D)*** IL-1β released from mouse BMDMs treated with ± 5 μM Miltefosine and primed with 1 μg/ml LPS, followed by transfection with 2μg of poly (dA-dT) for 3h to induce the AIM2 inflammasome. ***E)*** Cellular total cholesterol levels were determined in mouse BMDMs treated with ± 5 μM Miltefosine for 16h or 1 mM cyclodextrin for 2h. ***F)*** IL-1β released from mouse BMDMs treated ± 1 mM cyclodextrin for 2h or ± 5 μM Miltefosine for 16h followed by LPS and ATP treatment (mean ± SD, N = 3, different letters above the bars represent p<0.05 by ANOVA with Bonferroni posttest).

### Miltefosine reduced endotoxin mediated mitochondrial ROS production and loss of mitochondrial membrane potential

Mitochondrial ROS drives NLRP3 inflammasome assembly^39^ and mitochondrial cardiolipin provides a docking site for NLRP3 protein binding and inflammasome assembly^40^. As shown in ***Fig. 7A***, LPS treatment induced ROS production in control BMDMs as evident by increased staining with MitoSox Red while the Miltefosine pretreated cells showed decreased MitoSox staining upon LPS treatment. To quantify the MitoSox signal, we performed flow-cytometry and found that Miltefosine treated cells showed a significant 43% reduction in LPS-induced MitoSox staining (***Fig. 7B***). Next, we tested the effect of Miltefosine on the mitochondrial membrane potential. Control and Miltefosine pretreated BMDMs were incubated with ± LPS for 1h and stained with tetramethylrhodamine (TMRM) for 30 min at 37°C, followed by fluorescent microscopy of live cells. The cells treated with LPS showed marked reduction in TMRM staining while cells pretreated with Miltefosine partially rescued the mitochondrial membrane potential (***Fig. S10***). To quantify the TMRM signal, we performed flow-cytometry and found that LPS treated cells showed a significant 41% reduction in TMRM staining signal while cells pretreated with Miltefosine showed only a 24 % reduction (***Fig. 7C)***. Given that Miltefosine induced autophagy in macrophages, we tested role of Miltefosine in removing damaged mitochondria from via mitophagy. Live RAW264.7 macrophages were loaded with 500 nM mitophagy indicator dye for 30 min at 37°C, followed by treatment with ± Miltefosine for 4h. The control cells showed weak fluorescence while cells treated with Miltefosine showed bright fluorescence (***Fig. S11)***. To determine if the mitochondria were indeed fused with lysosomes, a lysosomal stain was used. As compared to control cells, the cells treated with Miltefosine showed markedly increased co-localization of the mitophagy and lysosome dyes (***Fig. 7D***). Thus, Miltefosine may inhibit inflammasome assembly by decreasing mitochondria ROS, preventing LPS mediated damage to mitochondria, and by removing damaged mitochondria through mitophagy.

**Figure 7.**
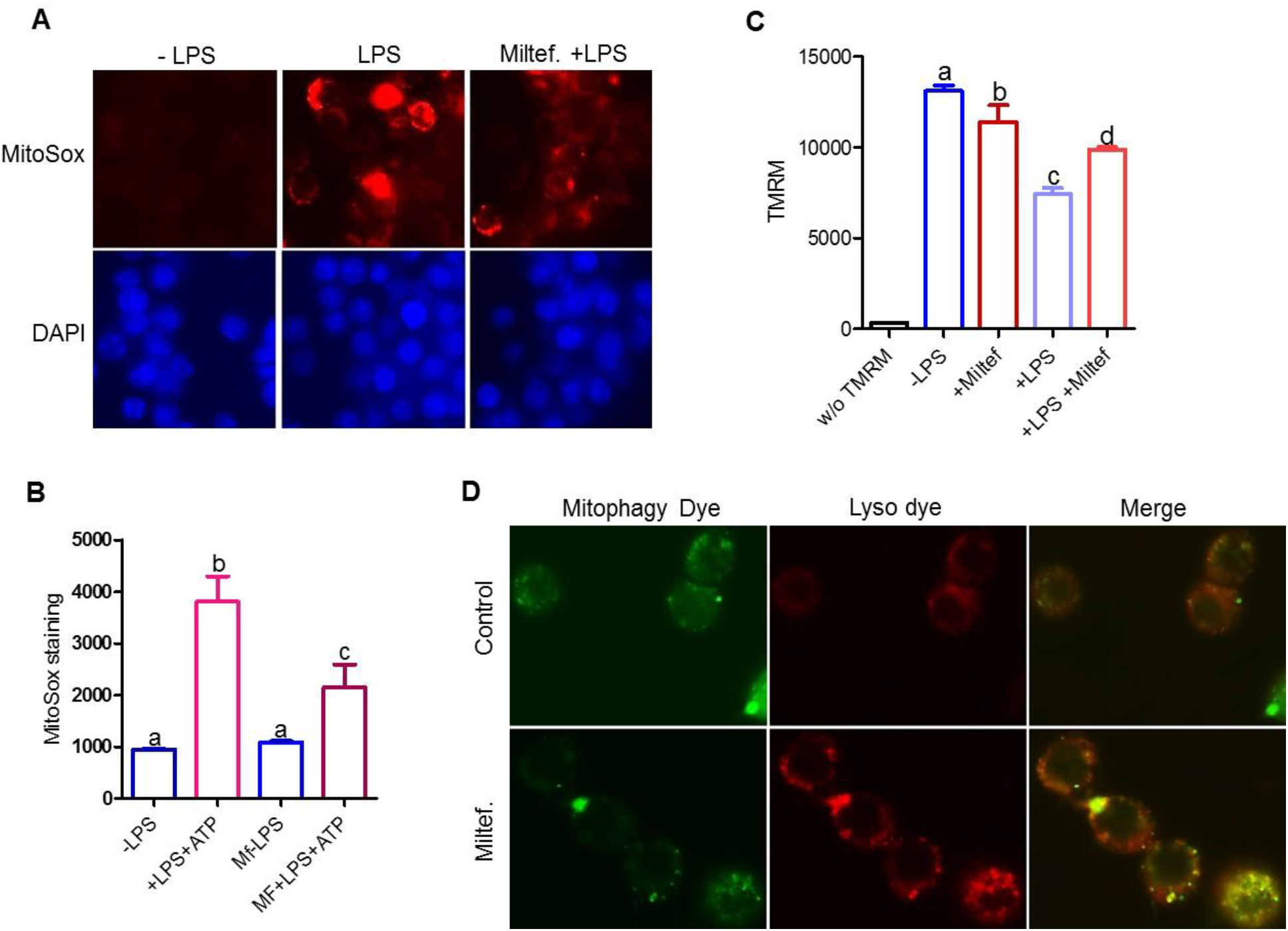
Miltefosine alters mitochondrial homeostasis and induces mitophagy. ***A)*** BMDMs pretreated ± 5 μM Miltefosine and treated ± 1 μg/ml LPS were stained with MitoSox to observe mitochondrial ROS in live cells. ***B)*** Flow-cytometry analysis showing quantification of MitoSox staining of live BMDMs treated as indicated (mean ± SD, N = 3, different letters above the bars represent p<0.01 by ANOVA with Bonferroni posttest). ***C)*** Live cell flow cytometry of mitochondrial membrane potential detected by TMRM staining of BMDMs pretreated ± 5 μM Miltefosine and treated ± 1 μg/ml LPS (mean ± SD, N = 3, different letters above the bars represent p<0.05 by ANOVA with Bonferroni posttest). ***D)*** RAW264.7 cells treated ± 7.5 μM Miltefosine and live cells were stained for mitophagy and lysosomes.

## Discussion

We found that Miltefosine has multiple activities in macrophages. Cholesterol release in the absence of an exogenous acceptor has been shown to be due to microparticle generation that is increased by ABCA1 expression^17^, and Miltefosine effects on the plasma membrane may alter membrane fluidity to increase microparticle generation. The Miltefosine increase in ABCA1-mediated cholesterol release requires functional lipid floppase activity as the ABCA1 mutant isoform defective in PS and PIP2 floppase activity showed lower cholesterol release to media in presence of Miltefosine. Addition of apoA1 to ABCA1 expressing cells in presence of Miltefosine did not showing additive effect, indicating that apoA1 may compete with pathways that are effluxing cholesterol in apoA1 independent manner. In addition Miltefosine delocalize PIP2 and we have shown before that apoA1 needs PIP2 for binding to cell-surface, thus apoA1 may have restricted access to ABCA1 in Miltefosine treated cells^15^. Thus, the Miltefosine mediated increase in cell surface PS and the disruption of lipid-rafts may lead to a redistribution of lipids to promote microparticle release.

PIP2 plays an important role in membrane ruffling, microvilli formation, endocytosis, phagocytosis and the attachment of membrane to cytoskeleton^41^. Miltefosine was initially identified as anti-cancer compound and is known to inhibit AKT signaling pathway^10^. Alkylphosphocholines such as Miltefosine are proposed to prevent plasma membrane recruitment of the PH domain of AKT by disrupting plasma membrane microdomains^42^. We speculate that Miltefosine effects on PIP2 localization could also play a role in AKT inhibition as reduced localization of PIP2 at plasma membrane by Miltefosine can deplete PIP3 pool at plasma membrane and reduce AKT signaling.

Autophagy plays a protective role in atherosclerosis and it is impaired in advanced human plaques^7,8,43^. Autophagy is controlled by multiple pathways responding to stimuli such as the status of cellular energy (AMP-dependent protein kinase, AMPK) or amino acid availability (target of rapamycin, TOR). Miltefosine induced basal autophagy in macrophages as evident by increased cytoplasmic p62 and LC3-GFP puncta. We showed that Miltefosine treatment increased the turnover of cholesterol esters in cholesterol loaded cells, supporting the notion that Miltefosine increased degradation of lipid-droplets via lipophagy. The increased phosphorylation of AMPK due to Miltefosine may lead to induced basal autophagy and mitophagy that we observed in Miltefosine treated cells.

Lipid-rafts play an essential role in recruiting TLRs to plasma membrane to initiate a signaling cascade that allows cytoplasmic NF-kB to shuttle inside the nucleus and upregulate inflammatory genes such as pro IL-1β mRNA and NLRP3. Miltefosine markedly reduced the LPS mediated induction of pro IL-1β mRNA. However, NLRP3 mRNA levels after LPS induction were similar in Miltefosine treated and control cells. Since both pro IL-1β and NLRP3 mRNAs are induced by NF-kB, the different response to NF-kB may in part be explained by their different degrees of induction, as NLRP3 mRNA was modestly increased (3-fold) compared to the robust (30,000-fold) induction of pro IL-1β mRNA. The differential regulation specificity for targets of NF-kB target have been described earlier^44–46^. There is the possibility that mRNAs of NF –kB targets, NLRP3 and IL-1β, are regulated differently at posttranscriptional level. The NLRP3 protein is stabilized via deubiquitination^34^, similarly there is possibility of NLRP3 mRNA is being stabilized while IL-1β mRNA is not. In addition, the synthesis of NLRP3 mRNA and IL-1β mRNAs, while both under control of NF-kB, may require different nuclear factors for transcription and Miltefosine treatment may affect these factors differentially. Thus, the detailed mechanism for the TLR-mediated induction of these two genes may differ.

The levels of NLRP3 inflammasome components were unaltered in control vs. Miltefosine treated cells. The intriguing question raised by these data is that if all inflammasome components are present and more or less equally expressed in control vs. Miltefosine treated macrophages, why is the NLRP3 inflammasome not assembled in Miltefosine treated cells? Recently it was shown that NLRP3 is localized to the trans golgi network via binding to PI4P prior to inflammasome assembly^47^. Miltefosine may work at this or other steps to block inflammasome assembly. Alternatively, Miltefosine may diminish K+ efflux which is required for activation of inflammasome or alter NLRP3 binding proteins such as NIMA-related kinase 7 (NEK7) that acts downstream of potassium efflux to regulate NLRP3 oligomerization and activation^48,49^. Another important factor regulating inflammasome function is mitochondria. Mitochondrial dysfunction or oxidized mitochondrial DNA fragments can also serve as signal for NLRP3 inflammasome activation^50,51^. Similar to our work, a recent paper from Karin lab showed inhibiting choline uptake leads to activation of AMPK and mitophagy and dampening of NLRP3 inflammasome activity and IL-1β release^52^. Miltefosine, a ephosphocholine analogue can inhibit phosphatidylcholine (PC) biosynthesis^53^, thus Miltefosine may be activating AMPK via PC depletion, leading to induced mitophagy and NLRP3 inflammasome inhibition.

## Material and Methods

A detailed description of the materials and methods used in this study is provided in *SI Materials and Methods*. Basic methods are summarized below.

### Cholesterol release assay

Cholesterol release assays were performed in HEK293-ABCA1-GFP cells, BHK cells, or RAW264.7 murine macrophages as described earlier^15^. Other details are described in *SI Materials and Methods*.

### Lipid-Raft quantification

The lipid-raft were visualized using Alexa647-labeled cholera toxin B subunit and fluorescent microscopy, with quantification by flow-cytometry assay as described before^25^. Other details are described in *SI Materials and Methods*.

### Total/FC measurements

RAW264.7 cells were loaded with 100 μg/ml AcLDL for 16 h at 37 °C with or without 2 μg/ml ACATi (Sandoz 53-035, Sigma) treatment for 2h. The total cholesterol and free cholesterol levels were determined by using enzymatic assay as described earlier^26^.

### Cell-surface PS and NBD-PS translocation

Cell-surface PS and translocation of NBD-PS were described earlier^25^. Other details are described in *SI Materials and Methods*.

### Western blotting

Details are described in *SI Materials and Methods*. Total and cell-surface ABCA1 levels were determined as described earlier^25^.

### PIP2 cellular reporter assay

RAW264.7 macrophage cell lines stably transfected with 2PH-PLCδ-GFP plasmid was described earlier^15^.

## Author Contributions

K.G. conceived, designed, performed and directed research. A.J.I., H.L., and J.H. performed research. A.J.I, H.L., J.H., H.A., J.D.S., and K.G. analyzed data. A.J.I, J.D.S, and K.G. drafted the manuscript. All authors critically reviewed the manuscript.

## Acknowledgments

This research was supported by American Heart Association scientist development grant SDG25710128 to K.G and NIH grants R01 HL128268 and R01 HL130085 to J.D.S.

